# Anonymized Somatic Tumor Twins (STTs) enable open genome data sharing and use in research and clinical oncology

**DOI:** 10.64898/2026.04.01.715771

**Authors:** Nicolás Gaitán, Rodrigo Martín, Daniel Tello, Elisa Benetti, Michela Riba, Luca Licata, Marc Arbonés, Romina Royo, David Olmos, Marco J. Morelli, Giovanni Tonnon, Elena Castro, David Torrents

**Author notes:** These authors contributed equally to this work (Nicolás Gaitán, Rodrigo Martín).

## Abstract

The study of somatic variants from tumor genomes is fundamental to cancer research and clinical decision-making. However, existing data protection frameworks impose restrictions on the use and sharing of these variants in conjunction with sensitive germline information. To overcome these challenges, we developed GenomeAnonymizer, the first method to anonymize short-read DNA sequences from tumor-normal pairs. This generates Somatic Tumor Twins (STTs), an anonymized version of the original data that preserves the donor’s privacy while retaining somatic tumor information and sequencing noise. This method successfully removed all detectable germline variants from the 47 PCAWG-Pilot samples. We further demonstrate that Whole-Genome Sequencing (WGS) STTs preserve more than 98% of the original somatic variants, enabling reliable downstream analysis that replicates somatic-related findings from the original samples, including cancer driver genes, mutational signatures, and intratumor heterogeneity. Importantly, we also show that STTs can reproduce the identification of actionable genes and downstream clinical interpretations and decision-making. We generated a cancer cohort of STTs matched with synthetic clinical data that could be openly shared and used across projects and centers worldwide. This paradigm-shifting approach will accelerate discovery and clinical translation in oncology and enable the robust benchmarking of genome analysis and large-scale data infrastructures.

## INTRODUCTION

Independent of the molecular background of cancer-initiating cells, most tumors form and progress through the acquisition of somatic mutations. Despite germline variants also playing a critical role in clinical oncology, particularly in cancer risk prediction and therapeutic decision-making, the systematic study of somatic variation has become a cornerstone of precision oncology, enabling the discovery of fundamental cancer driver genes and related pathocellular processes [1, 2]. These discoveries have improved tumor classifications, diagnoses, and treatments geared towards personalized oncology [3–5]. In this context, the collective impact of sharing, integrating and reusing somatic genomic data could further enhance biomarker discovery and its translation into clinical decision-making to a whole new level [2, 6], for example, by allowing the exchange and combination of multiple datasets from different projects and the analysis of primary care information that lack explicit consent for secondary use. In addition, the availability of real open somatic genomic data would also accelerate the development, standardization and benchmarking of data infrastructures for research and clinical oncology, where reliance on in-silico or fully synthetic datasets has so far limited translatability to real-world applications [7–9]. Despite these clear benefits, current data protection frameworks worldwide severely restrict the open sharing and use of tumor genome data. While somatic variants alone pose no identification risk [10–12], the presence of sensitive germline information renders tumor sequencing data identifiable [12], and thus subject to ethical and legal frameworks that protect individuals’ privacy [11].

Various strategies have been proposed to enable research while preserving donors’ privacy, including genomic beacons [13], and encryption-based methods, yet these approaches have proven vulnerable to re-identification attacks [14–16]. Given that the analysis of short-read whole genome sequencing provides an average of 4.5-5 million germline variants per individual [17, 18], partial anonymization protocols also remain insufficient due to a residual germline variation that could still allow for the reidentification of an individual [19]. Other efforts have focused on generating synthetic somatic mutation datasets using AI, which, while theoretically not vulnerable to reidentification, do not fully reproduce the biological properties of real tumor data [12, 20–23]. Nevertheless, the complete removal of germline variation from single-cell transcription data has been shown to enable the sharing of cell expression profiles while preserving patient anonymity [24], suggesting that a similar approach could be applied at the genomic level for tumor genomes while conserving the original somatic variant information.

To address these limitations and facilitate the open sharing of somatic tumor-normal whole-genome sequencing (WGS) datasets, we developed GenomeAnonymizer, a novel algorithm that generates Somatic Tumor Twins (STTs) by eliminating all germline variation from the original sequencing data. As a result, STTs preserve the original tumoral somatic mutation information, as well as read sequencing noise patterns. This approach enables researchers and clinicians to share and access tumor samples without risking patient confidentiality. Moreover, STTs can be tailored to specific research and benchmarking needs, enabling applications ranging from somatic variant discovery and interpretation to the evaluation and optimization of somatic analysis and data management protocols. Finally, we applied this method to the PCAWG-Pilot samples to generate a Somatic Cohort Twin that combines anonymized tumor-normal genome pairs with matching synthetic clinical data. This resource can be openly shared, under current data protection frameworks, and used for the design, development, and benchmarking of infrastructures for cancer data management and analysis.

## RESULTS

### Complete WGS anonymization: Removing all identifiable germline variation

As detailed in Figure 1, we developed an algorithm that anonymizes short-read sequencing data by removing germline variants and preserving tumor somatic mutations and sequencing noise. The method detects read alignment variation patterns concurrently shared between tumor and normal samples, and then replaces the altered read segments with the corresponding reference sequence (see Figure 1 and Methods). Considering that germline alleles may be masked by somatic Loss of Heterozygosity (LOH) in the tumor, recurrent read alterations detected only in the normal sample are also anonymized. These variation patterns are evaluated under stringent criteria (Figure 1) to ensure the detection and removal of all germline information while minimizing the risk of removing true somatic events and sequencing noise.

**Fig. 1:**
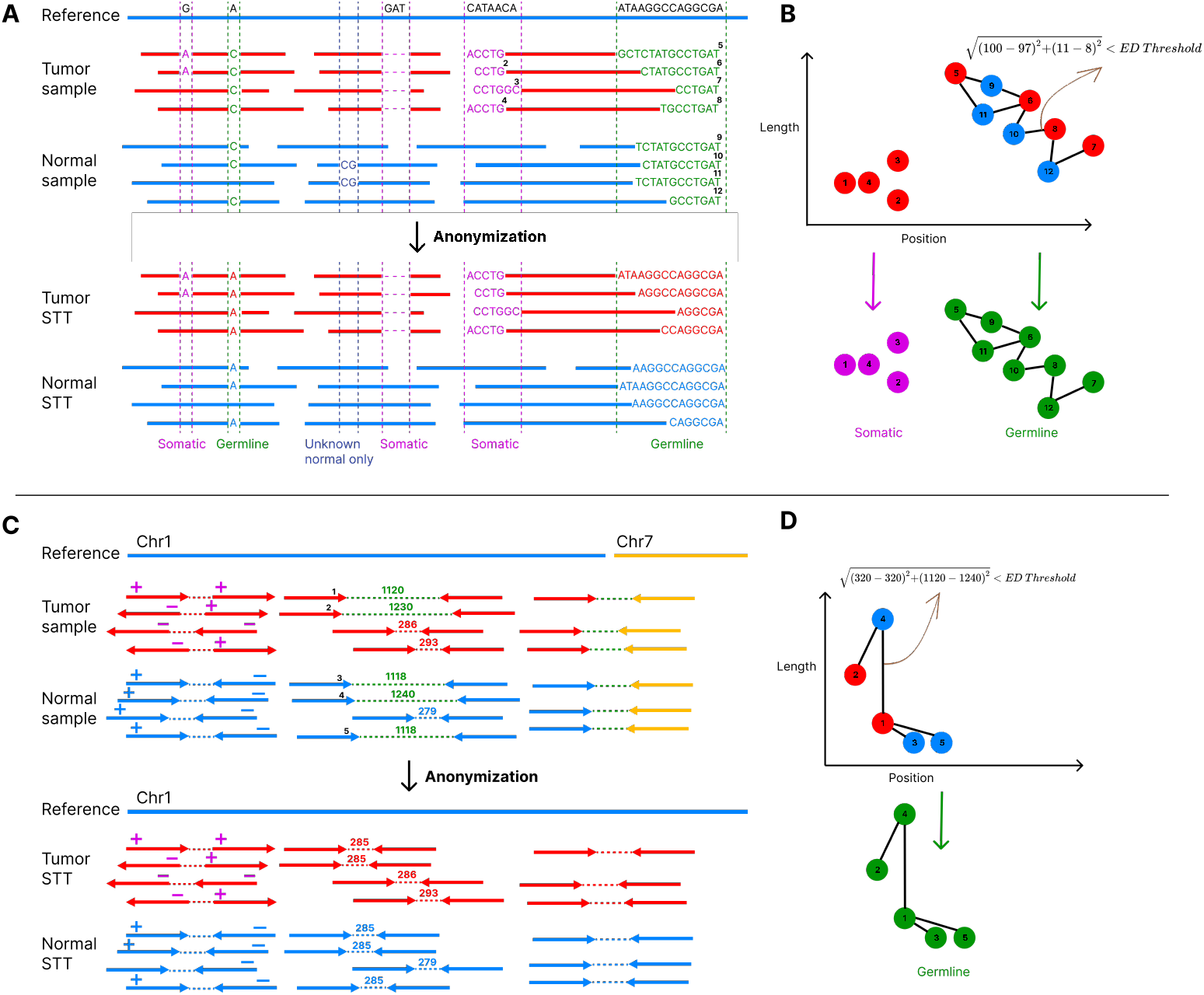
Schematic depiction of the anonymization algorithm. Systematically, the algorithm distinguishes both intra-alignment (mismatches, small insertions and deletions, and soft clipping) and inter-alignment variations [32] (discordant pairs or split alignments). Intra-alignment events are anonymized by restoring the reference allele within the affected reads, thereby removing germline SNVs, small insertions, and deletions. In contrast, inter-alignment events, which reflect Structural Variants (SVs), are corrected by replacing discordant read pairs with pairs generated from the reference genome, ensuring that mates map to the same chromosome with the expected average sample insert size and orientation between pairs. Together, these procedures eliminate germline variants of all types while maintaining the essential sequencing information for downstream tumor analyses. (a) showcases (left to right) how intra-alignment signals are characterized as somatic or germline, and how the latter are anonymized. Mismatches found in tumoral reads (e.g., G>A) are either treated as sequencing noise or classified as somatic variants in the STT. In contrast, equal alleles in both tumor and normal reads are classified as a germline SNV (A>C) and replaced with the reference nucleotide. The following insertion (CG) in two normal sample reads exemplifies a potential LOH event and is classified as a germline indel, thus eliminated from the reads. Two bases are added at either end of this modified read pair to maintain the original read length and insert size. (b) As the features of indels and clipping events often vary between read alignments, nearby signals are compared based on position and span, to assess if they come from the same germline variant. Tumor-only events unrelated to normal read variations are retained (pink nodes). In contrast, connected components indicative of a germline variant (green nodes) are anonymized by replacing their sequence in the read with reference bases. (c) Inter-alignment signals are characterized similarly, where discordant pair orientations, outlier insert size, or interchromosomal mates are grouped (d), if they are present in both tumoral and normal sample reads. To anonymize these signals, their orientation is corrected (first pair forward, second reverse) and/or the first pair is mated with a new one generated from the reference sequence, thus removing SV germline variation patterns. Additionally, unmapped reads and optical duplicates are removed from the STT.

To test this methodology, we applied it to the PCAWG-Pilot dataset from the PCAWG project [2], which includes 47 tumor-normal WGS pairs spanning 26 tumor types and sequencing depths ranging from 27x to 62x for normal samples and from 27x to 149x for tumor samples. On average, the anonymization algorithm modified 8.5% of each genome, leaving the remaining 92.5% intact. Then, to assess the degree of anonymization of the resulting STT samples, we used an inclusive strategy with 6 state-of-the-art variant callers for the detection of germline SNVs, indels, and SVs (see Methods), and compared the germline variant content of the original samples with their corresponding anonymized STTs. Across all 47 original sample pairs, a total of 628.7 million germline calls were identified (84.17% SNVs, 15.73% indels, and 0.95% SVs; approximately 3 million calls per sample per caller) (see Supplementary Table 1). Of these, only 23 calls (20 indels and 3 SVs) reappeared across 14 of the generated STT pairs. 21 of these calls were identified by a single caller, and one of them by two callers. A detailed inspection of each one of them showed, consistent with the behavior of the anonymization algorithm, that these spurious calls did not correspond to true germline variants but rather from imprecise alignments in repetitive regions (see Figures S1 to S21). Together, these results indicate a complete removal of germline variation within STTs.

### Evaluating the scientific value of STTs in tumor variant analysis

Having demonstrated the absence of detectable germline variants in STTs, we assessed both the quantity and quality of retained tumoral somatic information and their applicability across typical biomedical questions. Using a previously reported somatic variant identification pipeline [7], consisting of six callers that cover SNVs, small indels and structural variants (see Methods), we first evaluated and compared the number and characteristics of somatic variants before and after anonymization. From all experimentally validated somatic variants associated with the 47 original tumor samples, their corresponding STTs retained a median of 98±1.4%, 98±2% and 100±4.7% of the original SNVs, indels and SVs, respectively (Figure S22). In-depth analysis further revealed that a fraction of variants identified as somatic in the original samples but absent in STTs actually show evidence in both normal and tumor genomes, consistent with germline origin. These variants could be misinterpreted by variant callers in the original sample and are correctly removed during anonymization (Figure S23).

We next evaluated whether the fraction of somatic variation retained in STTs enables addressing fundamental questions in cancer research and clinical oncology. First, the high level of somatic variant retention allowed us to identify in STTs a median of 95% of the mutated coding genes detected in the original samples. When focusing on cancer driver genes [4], the median retention rate was 100%. We also assigned known COSMIC SBS96 [25] mutational signatures to both the original and STT samples (see Methods). Of all 47 original samples, 99% of signatures matched in their corresponding STTs (Figure S24). In addition, to assess whether the anonymization process affected the Variant Allele Frequency (VAF) of somatic variants retained in the tumor genome, and consequently downstream VAF-dependent analyses, we compared VAF distributions between original and STT samples and observed strong concordance (Figure S25). Finally, we evaluated whether clonal architectures could be defined from STTs with the same accuracy as the original samples. Because germline variants are required to accurately detect and correct somatic copy number variations (CNVs), which influence variant allele frequency distributions, somatic CNV analysis was performed exclusively on the original samples. We considered subclones that were supported by more than 5% of each of the samples’ variants and found that 46 out of 47 STTs showed an identical number of clusters compared to their original samples. Considering the entire cohort, 97% of variants were assigned to the same cluster in both original and STT samples (see Methods).

### STTs used in clinical tumor assessment

Having demonstrated the utility of STTs for somatic research, we also assessed whether STTs could be used in clinical settings. For this, we analyzed pathogenic mutations in key tumor biomarkers and compared recommended targeted treatments. We used the Clinical Genome Interpreter (CGI) [26] to translate the somatic information in each of the 45 PCAWG-Pilot WGS samples with known tumor type into clinical biomarkers. In all cases, the recommended treatments provided by CGI were identical between the original and the STT tumor-normal pairs for markers with Level A evidence [27], and 93% coincident when including markers with Level B evidence.

Although WGS is expected to be increasingly adopted by healthcare systems in the coming years [28], targeted gene panel sequencing is still a widely used technology for cancer genome analysis in oncological practice today. To evaluate whether GenomeAnalyzer can anonymize and retain value for gene panels, we generated an STT cohort from 50 tumor-normal prostate cancer paraphine-derived panels covering 128 genes (≈690x average depth) [29, 30]. Using the same strategy and consistent with the results obtained on WGS, gene panel STTs also showed a complete removal of detectable germline variants. Of 1.6 million total calls in the original samples, only one was detected in a single STT by a single variant caller, which also results from imprecise read alignments within a repetitive region. Next, we evaluated somatic information retention in gene panel STTs using three somatic variant calling approaches commonly used in clinical settings (see Methods). Analysis of biomarkers revealed notable variation across somatic pipelines, showing ranges of coincident CGI recommendations between 56 and 92% when considering level A evidence, and 52 and 92% when including level B evidence.

### Method limitations

Despite its advantages, our approach has the following limitations: (1) A primary and inevitable constraint is that STTs are unsuitable for pathogenic germline variant analysis, restricting the clinical applicability of STTs for the prediction of specific cancer risk, diagnosis, prognosis, treatment selection, and the determination of familial risk factors for certain tumor types based on specific genes (e.g., *BRCA, MMR, VHL*). (2) The method requires matched tumor-normal pairs to distinguish germline from somatic variants. (3) Low data quality can compromise the scientific value of STTs, especially for gene panels. Some somatic variants may be misclassified as germline and removed during anonymization due to tumor contamination. This challenge is already present in conventional analyses of hematological cancers [31]. (4) The anonymization algorithm was developed for short-read sequencing data and would require *ad hoc* adjustments for long-read sequencing. (5) Despite the removal of all detectable germline variants from gene panels, derived STTs should not be directly used for clinical decision-making without prior in-house benchmarking of somatic variant retention. (6) Finally, the strict removal of germline variants may also inadvertently eliminate sequencing noise or unclassified variations within unresolved repetitive regions of the genome.

### Anonymizing a complete tumor sample cohort

Finally, to address the increasing need for high-quality benchmarking data for developing and optimizing data infrastructures in cancer research and clinical settings, we generated a fully anonymized Somatic Tumor Twin cohort that falls outside the scope of applicable data protection legislation. This cohort is composed of 45 tumor-normal STTs derived from the PCAWG-Pilot dataset for which diagnosis information was available [2], and a corresponding newly synthesized clinical dataset that aligns with each tumor type and driver content (Figure 3). These Clinical data include basic demographics (gender, age) and key clinical information (tumor type, primary therapy, surgery). Combined with experimentally validated somatic variants in the PCAWG-Pilot dataset, this STT cohort provides a robust resource for benchmarking emerging infrastructures and solutions for the discovery, access, interoperability, and analysis of cancer genomic and clinical data across federated and centralized environments.

## DISCUSSION

This study presents the first method that fully anonymizes cancer genome sequencing data while retaining its scientific value for the analysis of tumoral somatic variants. We show that germline variation can be effectively removed from whole-genome sequences, eliminating the primary source of donor re-identification, and thus preventing inference attacks [14, 15]. By generating Somatic Tumor Twins (STTs) from tumor-normal WGS genome pairs, we retained 98% of original somatic variants intact and demonstrated that fundamental cancer genome analyses, including the identification of pathogenic somatic variants on driver genes, clinically actionable biomarkers, mutational signatures, and intratumor heterogeneity, can be reproduced with accuracy comparable to that of the original data. Based on our conservative algorithm to remove germline variants and on the results of our validation analysis on WGS and gene panels, we found no evidence of residual germline variants within the remaining pool of somatic variants (germline leakage). These results suggest that STTs represent a secure approach for sharing somatic genomic information across research institutions in cancer projects, avoiding the administrative and legal constraints currently required.

STTs could therefore be applied in a variety of research and clinical scenarios where data protection regulations currently limit the use of somatic genome data (Figure 2). In addition to facilitating driver gene discovery, STTs may enable institutions to openly exchange tumor genomic information, potentially facilitating the development of advanced somatic-based Artificial Intelligence (AI) models for predicting tumor progression and treatment response. Moreover, STTs could also provide a secure mechanism to make primary clinical genomic data available for research. Under current policies, primary sequencing data cannot be reused without the explicit patient consent, which is often unavailable, leaving a large fraction of valuable genomic data inaccessible for scientific investigation. Transforming these primary cancer genome data into STTs through anonymization would enable the safe reuse of somatic information for research while preserving the donor’s privacy. In addition, anonymized STTs could facilitate the secure transfer of somatic genomic data between hospitals and research institutions, both nationally and internationally.

**Fig. 2:**
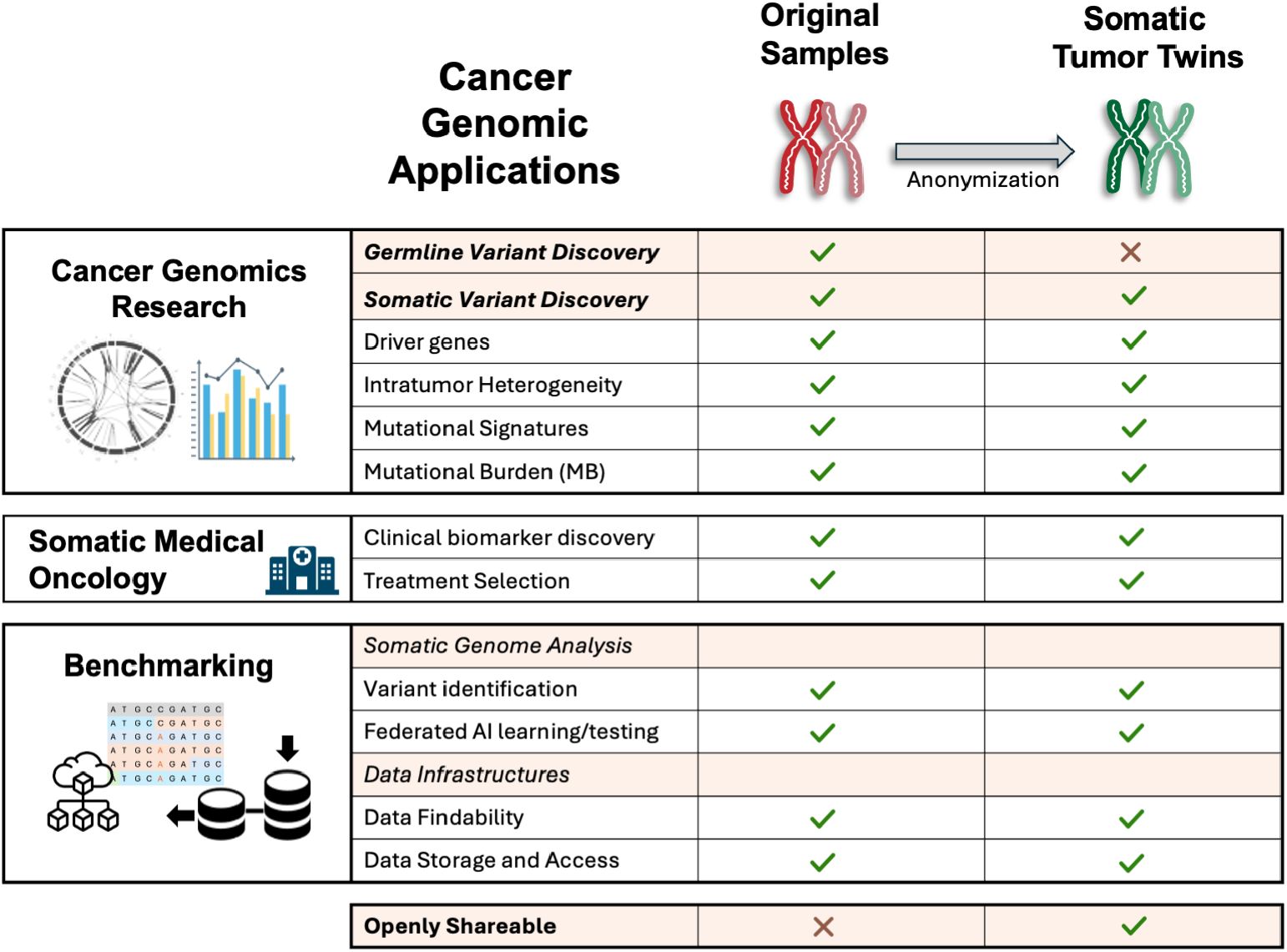
General overview of the different functionalities of Somatic Tumor Twins compared with original samples, including discovery, clinical application, and benchmarking of data infrastructures. A green checkmark indicates proven suitability for that particular functionality, whereas a red cross indicates a lack of suitability.

**Fig. 3:**
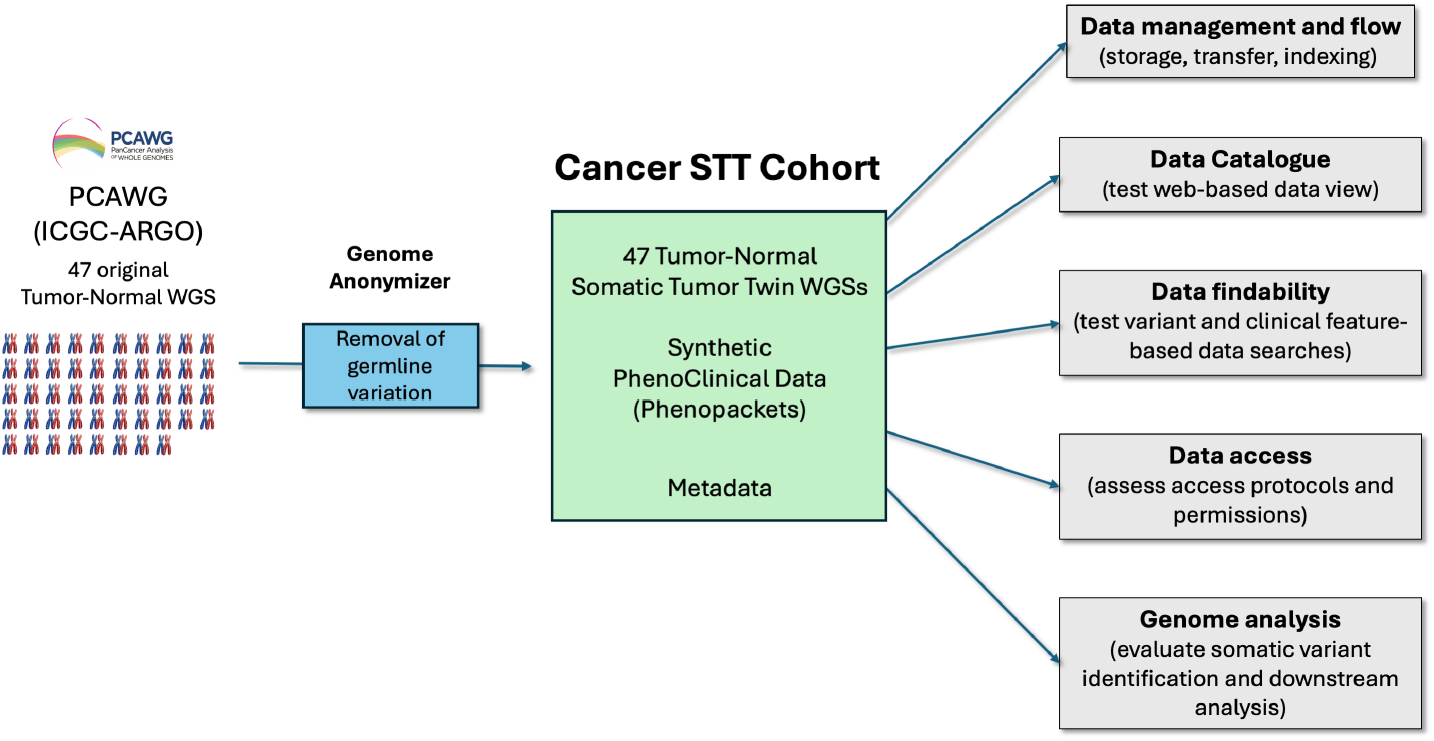
Generation and applications of a Somatic Tumor Twin cohort derived from the anonymization of the PCWAG-Pilot dataset. For each tumor-normal STT pair, somatic variant profiles were used to generate matching clinical annotations. The resulting dataset retains important genomic and biological characteristics of the original sample, while removing any personal information. The figure shows examples of scenarios of application, such as the development and calibration of data infrastructures and the benchmarking of somatic variant calling.

Finally, STTs can potentially be used in clinical settings for the discovery of actionable somatic biomarkers and thus to assess therapeutic options for personalized cancer treatments. Analysis of WGS STTs demonstrated complete FDA-approved biomarker retention (Level A) compared with original samples, whereas gene panel STTs vary according to the analysis pipeline, which invites further revision and benchmarking of somatic variant calling in clinical settings.

Overall, this study illustrates how privacy-preserving genomic resources can be derived from original sequencing data, enabling the open sharing of somatic tumor information data to enhance cross-institutional collaboration, somatic-based marker discovery, clinical decisions, as well as the development and benchmarking of data infrastructures. The remaining challenge is to adapt institutional policies and legal frameworks to enable the open exchange and reuse of STTs as a shared resource for cancer research and clinical translation.

## METHODS

### Algorithm description and implementation

To anonymize tumor and normal WGS samples, the STT algorithm detects and removes all sequence alignment discordances (signals) generated by germline variants from the reads mapped against the reference genome. Germline variants are identified as matching signals in at least one read from both the tumor and normal samples. This strategy derives from the conservative rationale that the same variant always generates similar signals and that a germline variant spawns at least one read signal in each tumor and normal sample. Moreover, as germline leakage may appear due to a somatic Loss Of Heterozygosity (LOH) in the tumoral sample, we also considered sequence discordances found in at least two normal sample reads as potential germline variants. Considering these scenarios also reduces the potential misclassification and removal of original sequencing errors and noise.

Each variant type, SNV, indel, or SV, can generate different mismatch patterns that the algorithm classifies into two main categories: those occurring inside the read-alignment (intra-alignment), or between read pairs or partial alignments (inter-alignment) [32]. Each of these signal types is identified and anonymized following different criteria and approaches.

First, intra-alignment signals are detected from the direct comparison of the read against the corresponding reference sequence, as well as from the CIGAR string of each alignment. For instance, mismatches with equal alleles and locus in multiple tumor and normal reads (or multiple normal reads) are considered evidence of an SNV and are replaced *in situ* with the corresponding reference nucleotide. Otherwise, because small insertions and deletions (indels), as well as clipped base pairs, exhibit higher variability, the algorithm measures their similarity by computing the Euclidean distance from their location and span to identify those that represent the same germline variant event. The pairs of signals that satisfy a lower Euclidean Distance than a variant type-specific threshold are considered as the same germline event, if at least one of them comes from the normal sample. Since indel and clipping signals are highly heterogeneous and somatic variants often appear close to the germline, an additional filtering step is applied to recover misclassified tumoral signals from anonymization. For this, hierarchical clustering is applied over groups of signals classified from the same event in order to generate precise clusters (in terms of signal position and length). If all of the signals in one of these clusters come from the tumor sample, they are considered somatic and thus excluded from removal. This fine-tuned algorithm minimizes the incorrect elimination of tumoral somatic mutations (see Figure 1a-b). The remaining germline signals are then anonymized according to the type of variation, by removing insertions from the read, adding nucleotides from the reference genome into the read across the span of deletions, or replacing clipped bases with the corresponding reference sequence instead (see Figure 1a-b).

In contrast, inter-alignment signals are identified from discordant read alignments, which can lead to the discovery of additional patterns, usually generated by SVs. The algorithm evaluates the similarity of inter-alignment signals, following the same procedure as intra-alignment signals, to find those that support the same germline variation event. These signals include inter-chromosomal pairs, abnormal read orientations, and/or anomalous insert sizes. In particular, anomalous insert size read pairs are identified as outliers (lower 5th and upper 95th percentiles) in the insert size distribution estimated from the original samples. Discordant read signals from pairs located on different chromosomes or outlier insert sizes are anonymized by eliminating one pair and generating its corresponding mate from the reference genome, such that their new distance satisfies the median of the sample’s insert size distribution. Abnormal orientations are corrected to ensure the leftmost pair maps on the forward direction (positive strand), and the rightmost pair on the reverse (negative strand) (see Figure 1c-d). On top of that, unmapped reads, optical duplicates, along with their pairs, are removed from the STT. Finally, to reduce spurious signals in abnormally high coverage regions (repetitive or low mapability loci), reads are included until a per-base depth threshold is reached (10,000 by default), after which, all other reads in this locus pileup are excluded along with their mates.

Due to the unique properties of gene panel datasets, we adjusted the algorithm to improve the retention of somatic information in STTs derived from them. Considering the deep sequencing depths of gene panels, the likelihood of a base differing from the reference, even in more than one read and simultaneously in the tumor-normal samples, is higher than in WGS. For that reason, two conditions were set to decide whether such an allele was anonymized or not. First, we set a hard threshold (1% of the total normal sample coverage at that position) to estimate the maximum amount of normal reads allowed to contain an allele before being anonymized. Second, if an allele is represented in more than one-third of the normal reads compared to the number of tumoral reads supporting the same allele, it is anonymized. These conditions ensure that somatic variants are not eliminated due to sequencing errors in the normal sample, while germline variants with low counts in both samples are correctly removed. Finally, as the allele frequencies of potential somatic variants in the STTs may be affected by the anonymization of some germline types (e.g., deletions) due to local depth changes, VAFs are estimated in the original tumor sample and are later used to correct tumor STT VAFs by replacing the reference base pair with the original allele, in as many reads as needed, until the original VAF is matched.

Lastly, we implemented this algorithm as an efficient, scalable, and self-contained software in GenomeAnonymizer, integrating multiple performance optimizations. A 74x tumor-normal pair can be anonymized under 5 hours using 16 cores and 24 GB of RAM, and a 140x pair in less than 9 hours under the same conditions. In addition, scaling core counts further decreases runtimes (1.24 average speedups from 16 to 32 cores) (see Figure S26). Among the most important factors for the computational efficiency of GenomeAnonymizer, we developed a pileup solution that provides reads on demand and simultaneously from a tumor-normal pair sample, returning reads only once on the first covered position, and then on nucleotides that differ from the reference. This allows a more efficient approach to pileup computation. We provide the code and the corresponding extendable classes for other potential applications outside GenomeAnonymizer.

### Validation of germline variants anonymization

We implemented a robust validation pipeline to ensure that the Somatic Tumor Twins (STTs) do not contain any identifiable germline variants (Figure S27). For this, we selected 6 germline variant callers, covering different algorithms, strategies, and scopes (SNV, indels, and SVs): Delly [33] (version 1.1.6), HaplotypeCaller (from GATK [34] 4.2.6.1), Strelka2 [35] (version 2.9.10), DeepVariant (version 1.6.1) [36], GRIDSS2 [37] (version 2.13.2) and Manta [38]. Each variant caller was used with default parameters and applied independently to the original and anonymized genomes, generating separate VCF files for each.

From the results of the calling phase, a variant was defined as germline if it was consistently identified in both the tumor and normal samples. We then used this collection of germline calls to test the anonymity of STTs by assessing whether these variants were fully removed. We used ONCOLINER [7] to carry out the intersection of germline variants between the original genomes and their STT counterparts. Comparison rules were tailored to variant types and sizes. In particular, SNVs and small indels were considered the same call if their chromosome, coordinate, and alternate allele exactly matched. Regarding SVs (insertions, deletions, and duplications with a length greater than 100bp, and inversions and translocations), two calls were considered the same if their breakends were located within a 100 bp window and their orientations were equal.

### Validation of somatic variant preservation in STTs

We conducted a comparative analysis using multiple variant callers on the original genomes to validate that somatic variant information remains in the anonymized Somatic Tumor Twins (STTs) (Figure S28). We built two different variant discovery pipelines (one for SNVs and indels that includes mutect2 (from GATK [34] 4.2.6.1), Strelka2 [35] (version 2.9.10), and SAGE [39] (version 3.0); and one for SVs that includes Delly [33] (version 1.1.6), Manta [38] (version 1.6.0), and GRIDSS2 [37] (version 2.13.2)), leveraging the optimal combination from PipelineDesigner [7].

Equivalent to the germline analysis, we used ONCOLINER [7] to characterize the intersection of somatic variant calls between the original genomes and their corresponding STTs across all variant-calling tools. Somatic variant preservation was quantified as the percentage of variants that were identical across the comparisons of all original samples and their STT versions, according to ONCOLINER’s analyses (following the same comparison principles described in the germline protocol).

### Evaluation of STTs’ value for tumor analysis

We performed multiple functional tumor assessment analyses to validate that the functional characteristics of the tumoral somatic mutation landscape were preserved in the STTs, including mutational signature detection and inference of intratumor heterogeneity. Additionally, we validated that variants affecting protein-coding and driver genes were kept in the STT.

To carry out mutational signature detection, we employed SigProfilerAssignment [40] to assign previously known mutational signatures to individual samples and 10 individual somatic mutations. We employed the COSMIC SBS96 catalogue and filtered out all signatures marked as unknown or sequencing artifacts [25], leaving 43 total signatures to analyze. Using the SNVs detected by the somatic variant calling pipeline, we considered that a mutational signature was preserved if SigProfilerAssignment showed presence or absence (either a non-zero or zero value, respectively) for that specific signature on the sample and its corresponding STT.

Regarding intratumor heterogeneity inference, we first extracted all somatic CNVs using `cnv_facets` (https://github.com/dariober/cnvfacets), which detects somatic CNV in tumour-normal samples [41]. This process was performed in the original samples, not the STTs, as it requires the germline information to be present. Then, we calculated the VAFs (by directly analyzing the alignment file at the corresponding loci) of all SNVs detected by the somatic SNV variant calling pipeline in both the original samples and their corresponding STTs. Finally, combining the somatic CNV information with the VAFs, we employed PyClone-VI [42] to infer the subclonal architecture of the tumors in both the original samples and STTs. We then filtered out all clusters that contained less than 5% of the total variants for that sample. Finally, using the remaining clusters, we counted the number of clusters and checked if the intersecting variants were in the same cluster in the original sample and its corresponding STT.

To assess whether the somatic variants that affect functional elements are conserved in the STTs, we used VEP [43] to annotate SNVs and indels, filtering out non-coding elements. Since VEP could not process structural variants due to their VCF representation in the original cohort, we developed custom scripts based on VariantExtractor [7]. Specifically, if a protein-coding gene intersected the span of an SV, then it was considered affected by this variant. For inversions and translocations in particular, only the genes that intersected their breakpoints were considered as affected, as these events usually disrupt the open reading frame of the gene. Using the same rationale, we also assessed the preservation of variants affecting cancer driver genes across SNVs, indels, and SVs. The source for the annotation of protein-coding genes to SVs was the Ensembl GHRCh37 transcriptome (v87) [44], and the cancer driver genes were collected from the IntOGen Catalog (release 2023.05.31) [4].

### Evaluation of STTs for clinical practice

To assess the applicability of Somatic Tumor Twins (STTs) in a real-world clinical sequencing context, we applied the anonymization algorithm to a prostate cancer cohort derived from the GENPRO12 study, which investigated the prevalence of prostate cancer related gene alterations and the impact in outcomes within the prostate cancer patients treated at Hospital Universitario 12 de Octubre, Madrid, Spain. GENPRO12 is part of the PROCURE Biomarkers Platform [29, 30].

For the present study, a subset of 50 prostate cancer tumor-normal pairs with available high-quality genomic data was selected for anonymization and downstream analysis. All patients were provided informed consent for somatic and germline genomic analyses, supported by the availability of peripheral blood samples for germline DNA extraction and formalin-fixed paraffin-embedded (FFPE) tumor tissue for somatic DNA analysis. FFPE samples underwent central pathology review to confirm tumor content and suitability for genomic profiling.

Tumor and matched normal samples were sequenced using a targeted hybridization-capture gene panel enriched for clinically relevant prostate cancer genes, including homologous recombination repair (HRR) pathway genes and other actionable cancer drivers. Sequencing was performed on Illumina platforms following standard protocols as described before [29, 30].

Reads were aligned to the GRCh38 reference genome using BWA-MEM (version 0.7.17), followed by duplicate marking and base quality score recalibration using Picard (version 2.7.1) and GATK (version 4.2.6.1). Somatic variant calling was performed using GATK’s Mutect2, SAGE [39], and Strelka2 [35]. These variant calls were used to evaluate the impact of anonymization on clinically relevant somatic alterations compared to the original samples.

### Clinical data generation

Clinical data have been structured into phenopacket-schema version 2 [45] starting from information available in the original manuscript of the PCAWG-Pilot cohort [2] and curating the addition of standard codes and structured definitions based on standardised oncology resources, based on the guide of the Minimal Dataset for Cancer [46]. Therapies have been referred where possible to HemOnc.org, the largest freely available wiki of oncology regimens and medications [47]. Surgical procedures have been coded with SNOMED vocabulary [48].

### Execution environment

All experiments conducted on the PCAWG-Pilot cohort were executed on a High-Performance Computing (HPC) environment composed of Intel Sapphire Rapids nodes (4th-Generation Intel Xeon Scalable Processors). All analyses performed on the tumor gene panels were carried out on an HPC environment composed of Intel Xeon Gold and AMD EPYC processor-based nodes with high-memory configurations.

## Resource availability

GenomeAnonymizer is publicly available as a container on DockerHub, with the following project ID and tag: ngaitan55/genomeanonymizer:latest. Additionally, its life development and source code are deposited on GitHub, as well as detailed installation and instructions at: https://github.com/Computational-Genomics-BSC/GenomeAnonymizer. The GFF3 file used for gene annotation on the GRCh37 reference can be found in the Ensembl FTP archive at https://ftp.ensembl.org/pub/grch37/current/gff3/homo_sapiens/Homo_sapiens.GRCh37.87.gff3.gz. The release 2023.05.31 of the compendium of cancer driver genes from Intogen is available on their website at https://www.intogen.org/download. PCAWG-Pilot dataset is available at https://docs.icgc-argo.org/docs/data-access/icgc-25k-data. The STT cohort, along with the synthetic clinical data phenopackets, is available at XXX (under submission). Variant caller containers and scripts are available at https://github.com/Computational-Genomics-BSC/somatic-variant-callers and https://github.com/Computational-Genomics-BSC/germline-variant-callers. The COSMIC mutational single-base substitution (SBS96) signature classification can be accessed at https://cancer.sanger.ac.uk/signatures/sbs.

## Author contributions

D.T. conceptualized, designed, and directed this work. D.T., G.T., M.M., D.O., and E.C. were principal investigators and contributed to supervising the study. N.G. and R.M. formalized, designed, and implemented the algorithm and software solution. N.G., R.M., D.Te., E.B., R.R., M.R., L.L., and M.A. contributed to the study methodology and data analysis. E.C. and D.O. contributed data to the study. The manuscript was written and compiled by D.T., N.G., R.M., D.Te. All authors were responsible for the preparation of the manuscript and the decision to submit for publication

## Acknowledgments

BSC-CNS discloses support for the research of this work from the European Union’s Horizon 2020 research and innovation programme under EUCANCan (grant agreement no. 825835), the Departament de Recerca i Universitats de la Generalitat de Catalunya (code: 2021 SGR 01626), the Science and Innovation Spanish Ministry under project BenchSV (PID2020-119797RB-I00/AEI/10.13039/501100011033), the Instituto de Salud Carlos III (ISCIII) project PMP21/00015 (PREGENLIF), co-funded by the European Union – NextGenerationEU (Recovery and Resilience Mechanism), and part of the project I+D+i PID2023-15867NB-I00 (PREVDIS), funded by MICIU/AEI/10.13039/501100011033 and by “ERDF/EU”. This work has been funded by the FPU23/03413 (Formación de Profesorado Universitario) scholarship granted to Nicolás Gaitán by the Spanish Ministry of Science, Innovation, and Universities. The PROCURE network, which provided data and samples for this study through the GENPRO12 cohort, was supported by CRIS Excellence 19-26, ‘Fundación Científica de la Asociación Española Contra el Cáncer (AECC)’ PROYE19045OLMO, and ‘Instituto de Salud Carlos III’ grants through projects PI13/01287, PI16/01565, PI19/01380 and PI22/01593 to DO, as well as PI19/01475 to EC. Additional support was provided by unrestricted grants from ‘Fundación CRIS Contra el Cáncer’ to DO (2012-2018) and to EC (2019-2022) and ‘Sociedad Española de Oncología Médica (SEOM)’ research projects to DO (2014) and to EC (2018, 2019). The Prostate Cancer Translational Research Unit of Hospital Universitario 12 de Octubre is supported by ‘Fundación FERO’.

All data used in this study were processed in accordance with applicable ethical guidelines and data protection regulations.

