## Supplemental Information for "Anonymized Somatic Tumor Twins (STTs) enable open genome data sharing and use in research and clinical oncology"

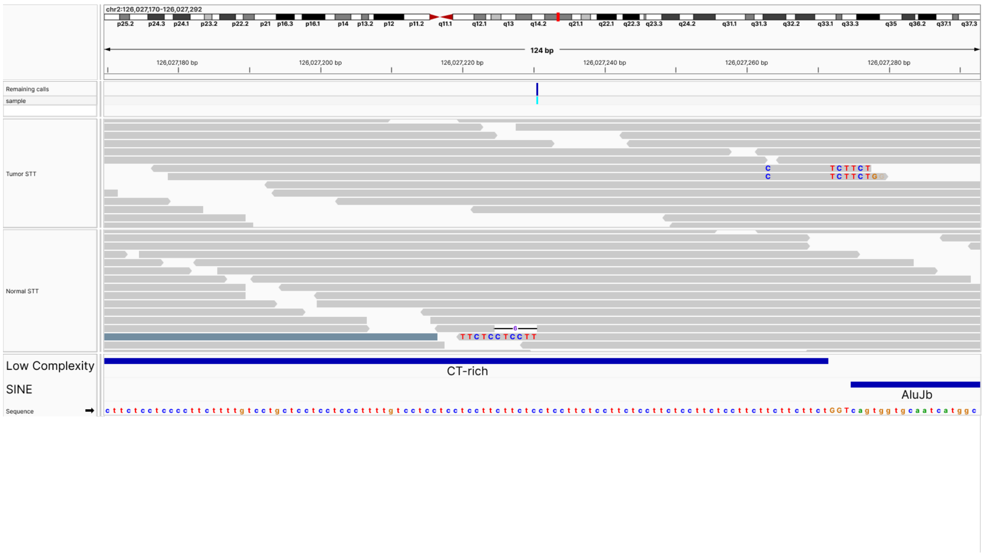


**Fig. S1**: **False positive call (1)**. False positive insertion call predicted by DeepVariant, likely due to ambiguous read-alignments in this low-complexity repetitive region, followed by an Alu element. Due to the presence of these repetitive elements, some alignment-generated variability appears in some read-alignments, which is taken as evidence of a variant by DeepVariant, even though it does not strongly support such an event in this scenario.

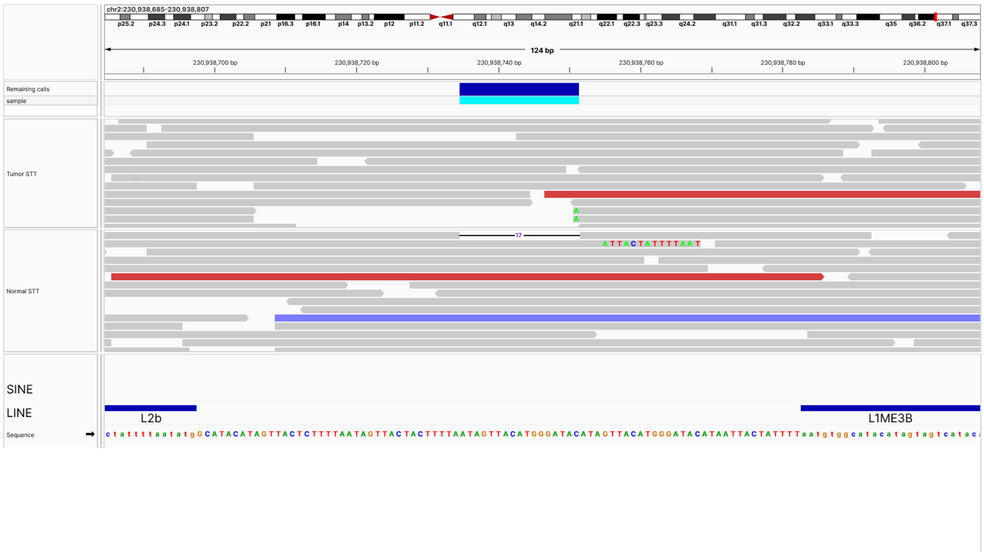


**Fig. S2**: **False positive call (2)**. False positive deletion call predicted by DeepVariant, likely due to ambiguous read-alignments in LINE-enriched repetitive region (Long Interspersed Nuclear Elements), as can be appreciated in the figure displaying the call flanked by two repetitive elements.

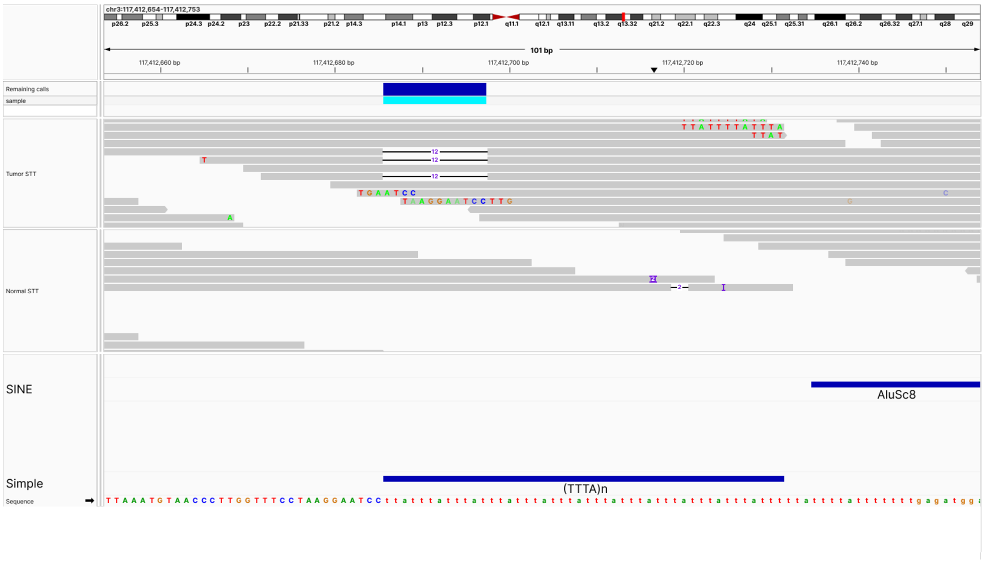


**Fig. S3**: **False positive call (3)**. False positive deletion call predicted by HaplotypeCaller, likely due to ambiguous read-alignments in this simple repetitive region, followed by an Alu element. In addition, although HaplotypeCaller likely mistook them as germline evidence, the Tumor STT contains read alignments that appear to come from a somatic variant; hence, GenomeAnonymizer did not modify them.

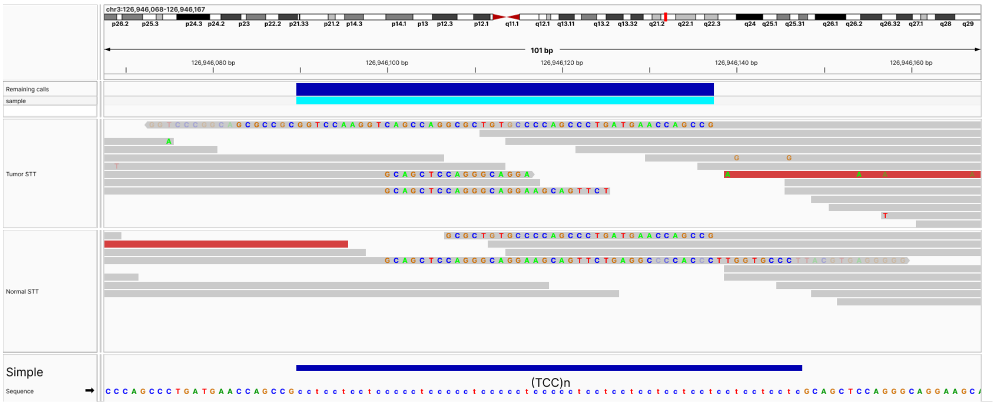


**Fig. S4**: **False positive call (4)**. False positive deletion call predicted by HaplotypeCaller, likely due to ambiguous read-alignments in this simple repetitive region (TCC repeat), evidenced by uneven soft clipping from the mapped reads shown in the figure.

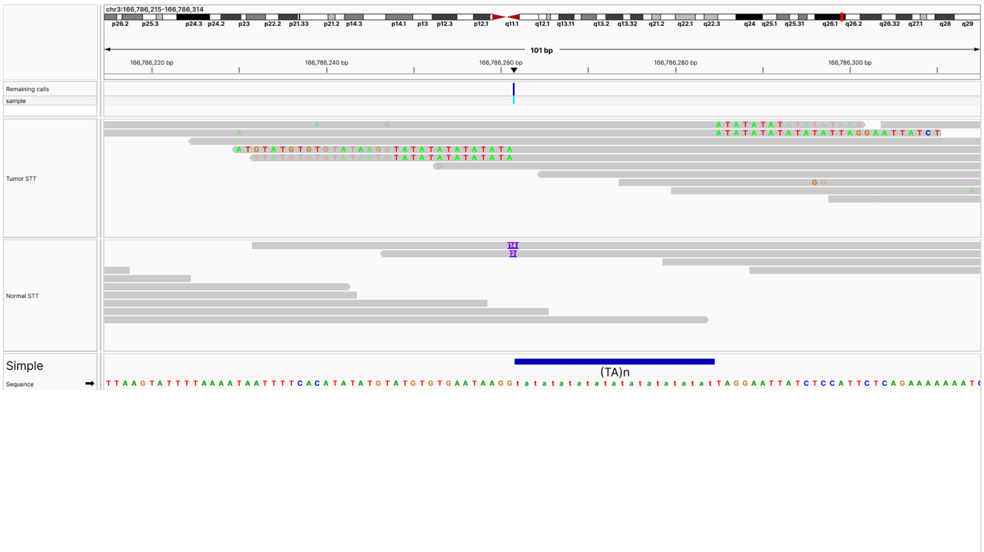


**Fig. S5**: **False positive call (5)**. False positive insertion call predicted by HaplotypeCaller, likely due to ambiguous read-alignments caused by a simple repetition (TA). These repetitive sequences are observed in the tumor reads that contain soft clip elements, whereas in the normal sample, they are aligned inside the reads. Although not annotated, additional simple repetitions can be observed in the reference genome, such as TG, which further reduces the precision of the mapping and thereby the accuracy of variant calling.

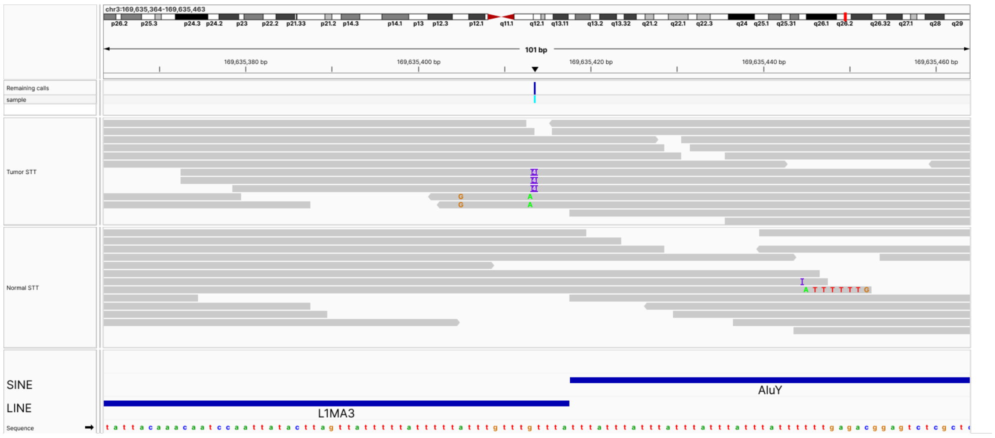


**Fig. S6**: **False positive call (6)**. False positive insertion call predicted by HaplotypeCaller, likely due to ambiguous read-alignments in a repetitive region, composed of adjacent LINE and SINE (Short Interspersed Nuclear Elements) elements. As shown in the figure, this variant call comes from distinct read alignment patterns, where the tumoral reads actually reveal a potential tumoral insertion. In contrast, the normal reads indicate variation from misalignment.

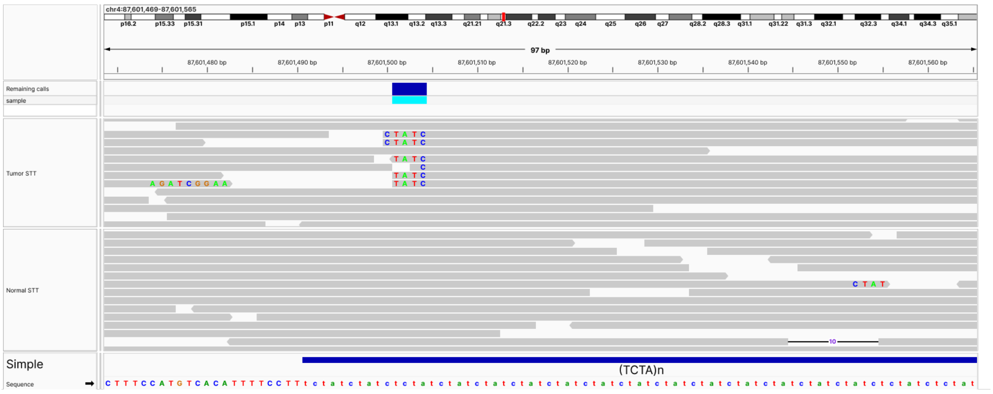


**Fig. S7**: **False positive call (7)**. False positive deletion call predicted by DeepVariant, likely due to ambiguous read-alignments in this simple repetitive region (TCTA). The alignment figure shows reads in both the tumor and normal STTs with different patterns, including softclips and even one read with a deletion, with differing lengths and coordinates.

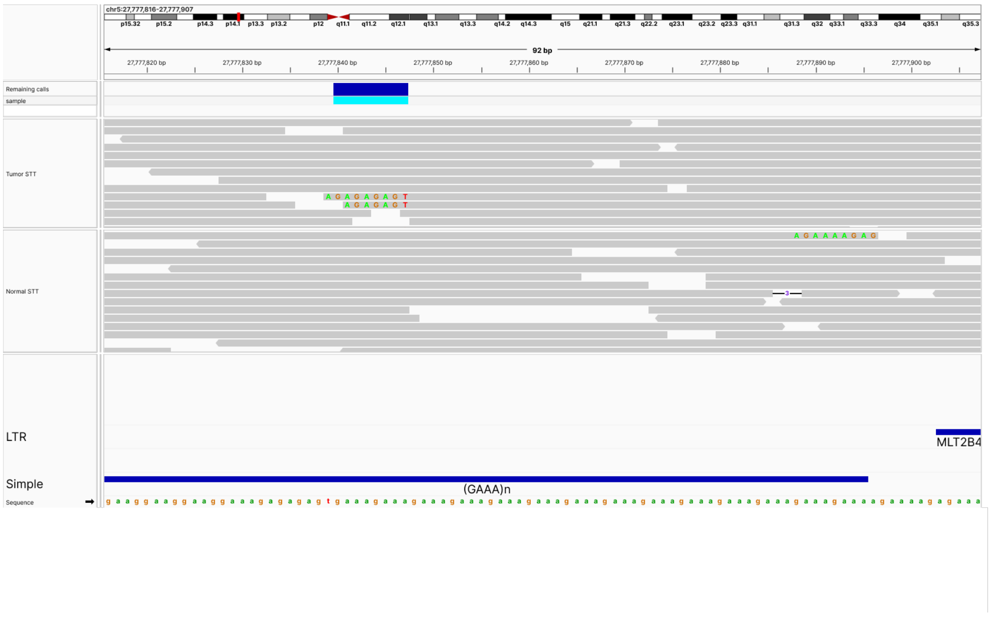


**Fig. S8**: **False positive call (8)**. False positive deletion call predicted by DeepVariant, likely due to ambiguous read-alignments in this simple repetitive region (GAAA). As seen in the displayed alignments, only the normal sample shows a read with a small gap, while some tumor reads contain softclips at different coordinates throughout the span of the reference repeat.

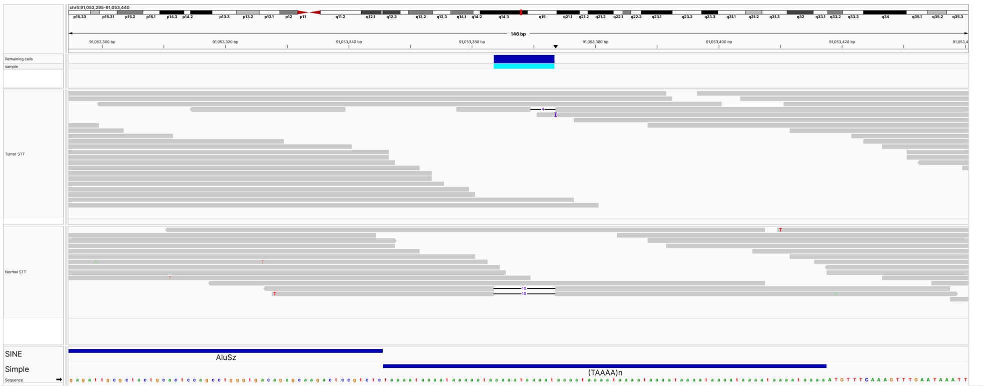


**Fig. S9**: **False positive call (9)**. False positive deletion call predicted by HaplotypeCaller, likely due to ambiguous read-alignments in this simple repetitive region (TAAAA), preceded by an Alu element. The alignment variation in repetitive regions generates spurious discordant gaps between reads, as shown in the figure.

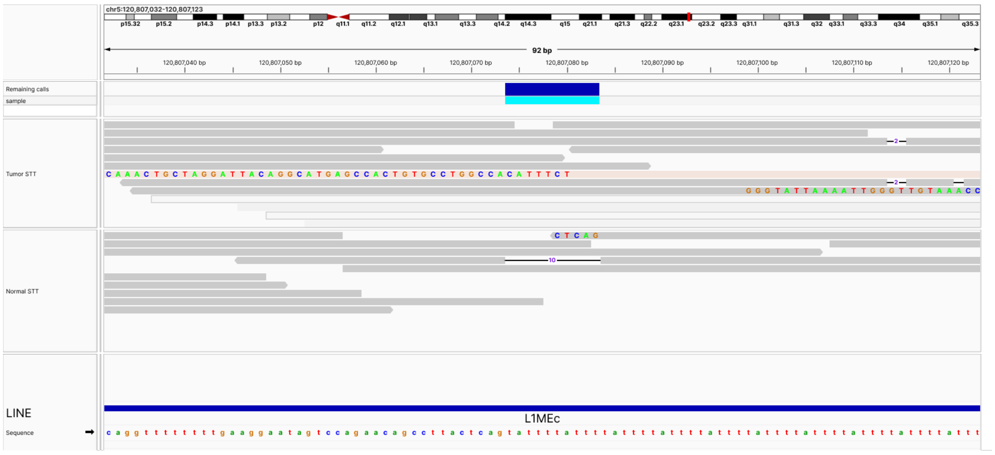


**Fig. S10**: **False positive call (10)**. False positive deletion call predicted by HaplotypeCaller, likely due to ambiguous read-alignments across this LINE repetitive element. It is also apparent that the evidence comes from a single deletion in one read and soft clips in other reads with different positions and lengths.

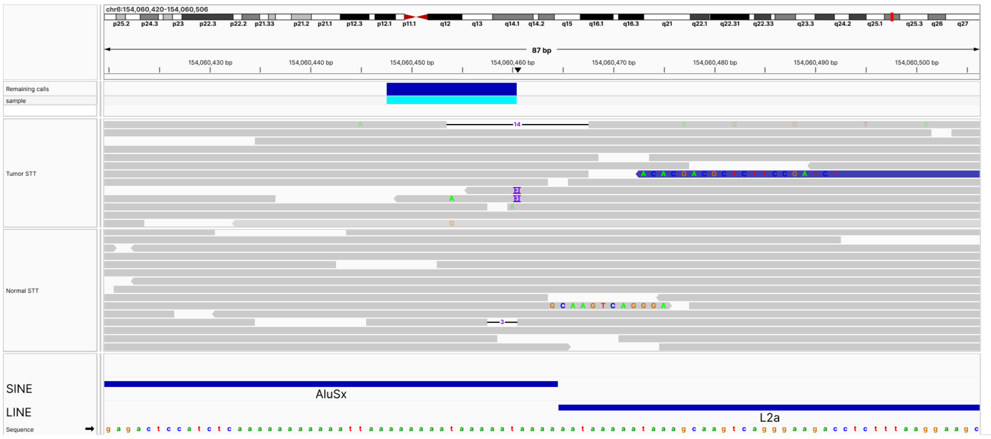


**Fig. S11**: **False positive call (11)**. False positive deletion call predicted by DeepVariant, likely due to ambiguous read-alignments in this repetitive region, comprising adjacent SINE and LINE elements. It can be appreciated how alignments are affected by the repetitions, generating apparently random insertions, deletions, and soft clips in multiple reads. Nevertheless, DeepVariant produces a deletion call, even though there is no variation consensus across read alignments.

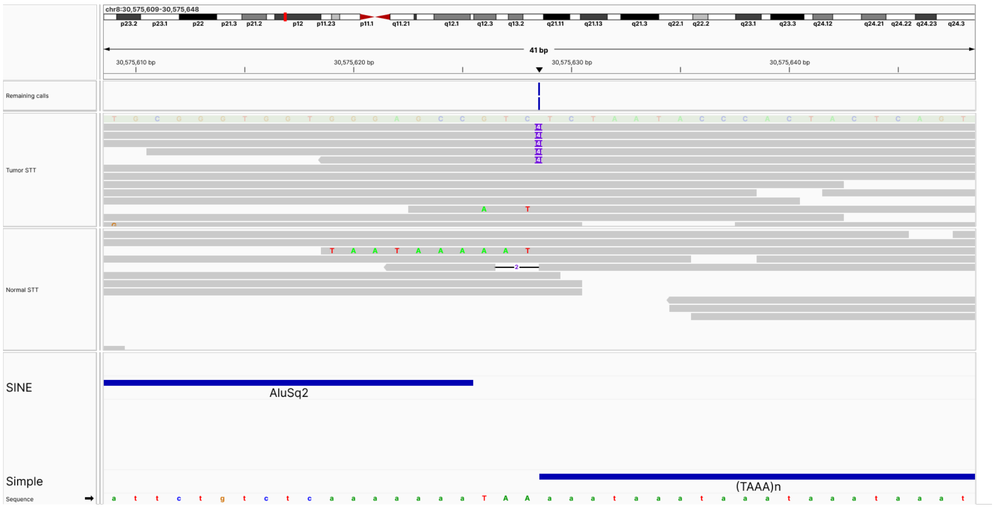


**Fig. S12**: **False positive call (12)**. This image displays the only call made by two tools (HaplotypeCaller and DeepVariant), consisting of a false positive insertion. In this case, the evidence from the tumoral read-alignments indicates a somatic insertion, while the normal sample contains a very different pattern, comprising a soft-clip and a deletion. While the two software agreed on this call, the weak supporting evidence, the difference in variation pattern between the two samples, and its presence in a repetitive region (composed of a SINE element and an adjacent simple repetition) do not support the presence of a true germline variant call.

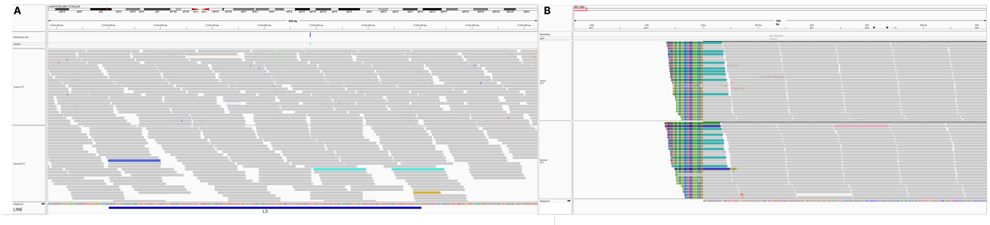


**Fig. S13**: **False positive call (13)**. False positive translocation call predicted by GRIDSS. In this case, a LINE repetitive element hinders the correct alignment of reads in the region shown in the left panel. Thus, supplementary alignments are generated (left panel), and partially aligned pairs are poorly mapped at the beginning of the mitochondrial chromosome (right panel). GRIDSS interprets this event as a translocation, even though the alignments are not reliable, which indicates a false positive event.

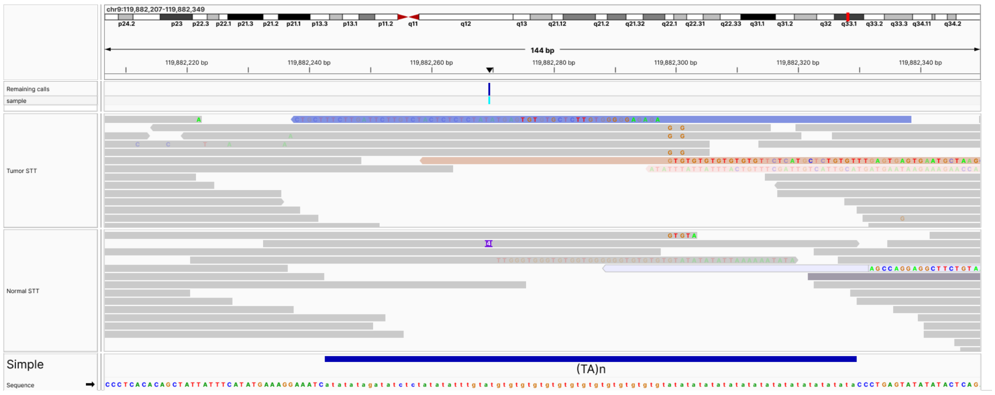


**Fig. S14**: **False positive call (14)**. False positive insertion call predicted by HaplotypeCaller, likely due to ambiguous read-alignments in a simple repetitive region (TA). In this case, it was apparently generated from soft-clips and one insertion, which, given their variability, do not constitute strong evidence for a true variant call, in addition to a low coverage in this region.

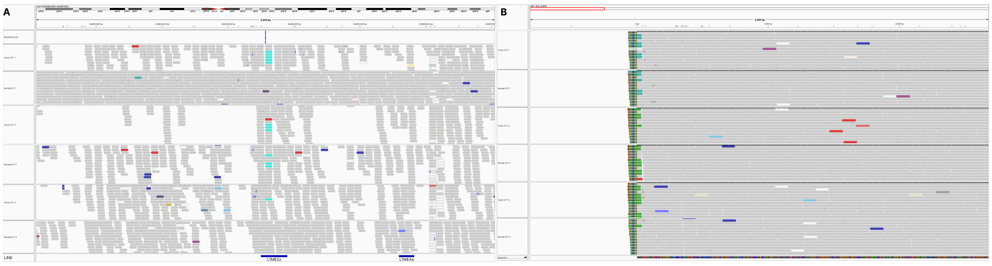


**Fig. S15**: **False positive call (15)**. False positive translocation call predicted by GRIDSS. In this case, two nearby LINE repetitive elements hinder the correct alignment of reads in the region shown in the left panel. Thus, supplementary alignments are generated (left panel), and partially aligned pairs are poorly mapped at the beginning of the mitochondrial chromosome (right panel). GRIDSS interprets this event as a translocation, even though the alignments are not reliable, which indicates a false positive event.

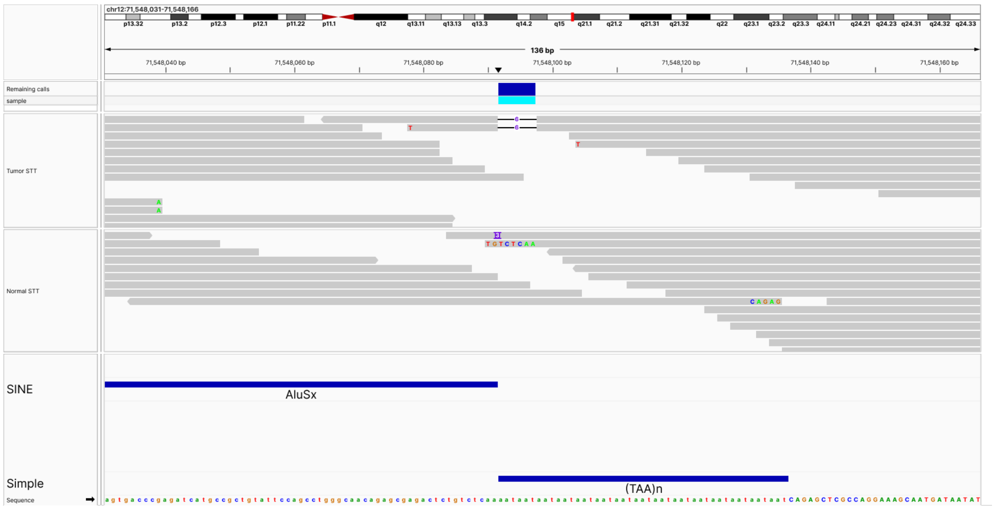


**Fig. S16**: **False positive call (16)**. False positive deletion call predicted by HaplotypeCaller, likely due to ambiguous read-alignments in this repetitive region, comprising an adjacent SINE element and a simple repeat (TAA). It can be appreciated how, in the tumor sample, there are two reads containing a deletion, while in the normal sample, a few reads contain soft-clips and an insertion. This variable alignment pattern indicates that the call was generated with conflicting evidence, pointing to a false-positive variant call.

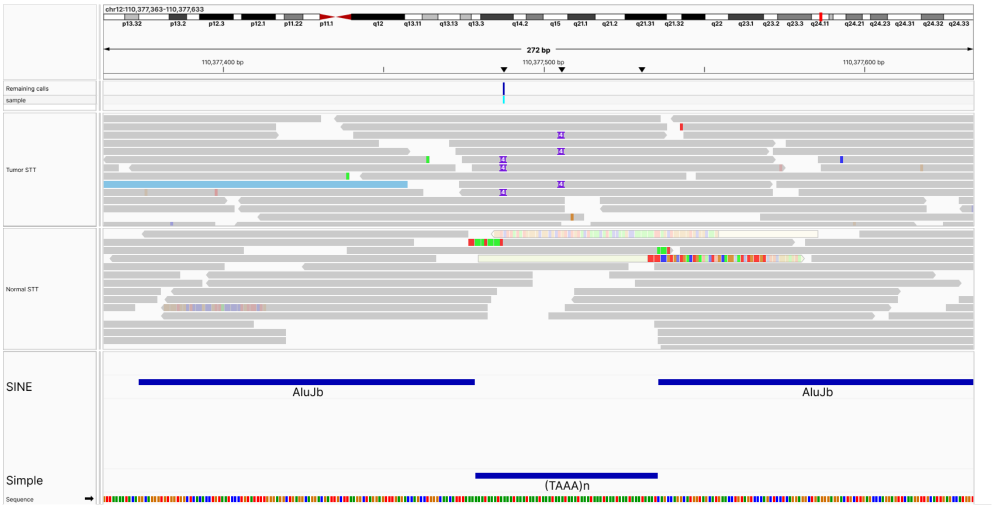


**Fig. S17**: **False positive call (17)**. False positive insertion call predicted by HaplotypeCaller. This figure displays a complex repetitive region comprising three elements in the following order: AluJb (SINE) - Simple repeat (TAAA) - AluJb (SINE). This complex pattern makes unambiguous alignment impossible, hampering variant calling quality. This is shown by the variability in both the tumor and normal samples, where reads in the tumor samples display either intraalignment insertions or soft-clips, respectively.

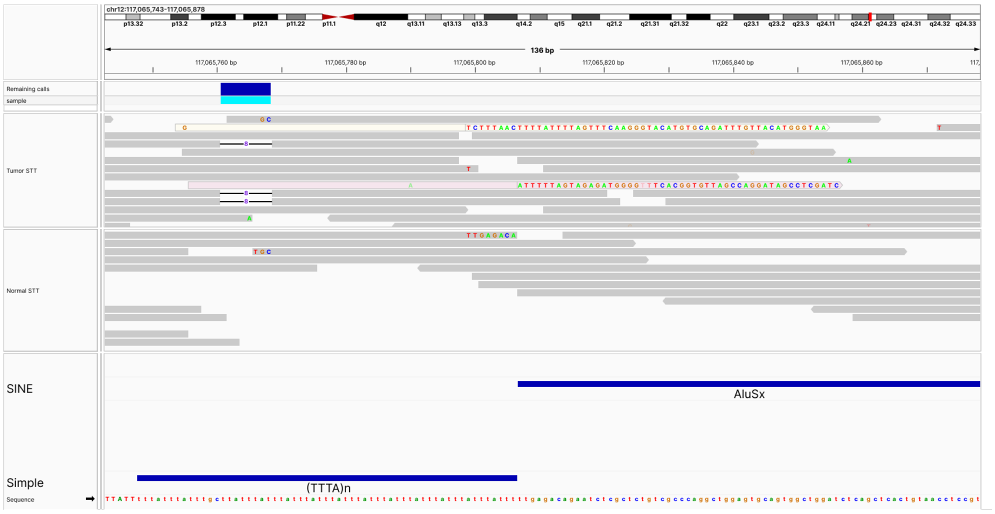


**Fig. S18**: **False positive call (18)**. False positive deletion call predicted by HaplotypeCaller, likely due to ambiguous read-alignments in this simple repeat region, followed by an Alu element. Due to the presence of these repetitive elements, some reads are aligned with gaps, while others contain unmapped sequences as soft-clip events. There is not enough evidence to consider it a true variation event instead of a mapping artifact.

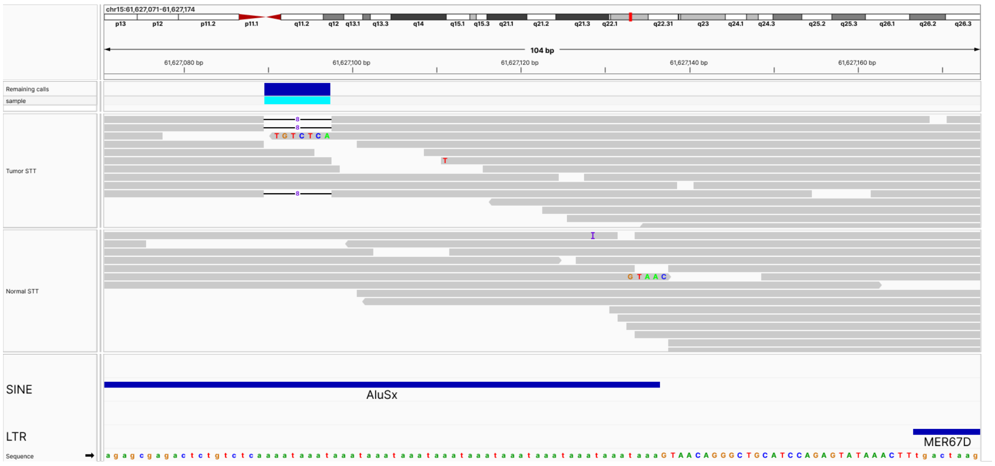


**Fig. S19**: **False positive call (19)**. False positive deletion call predicted by Strelka, likely due to ambiguous read-alignments in this SINE repeat region, followed by an LTR (Long Terminal Repeat). Due to the presence of these repetitive elements, some reads are aligned with gaps, while others contain unmapped sequences as soft-clip events. In this case, in the normal sample, there is no evidence of the variant.

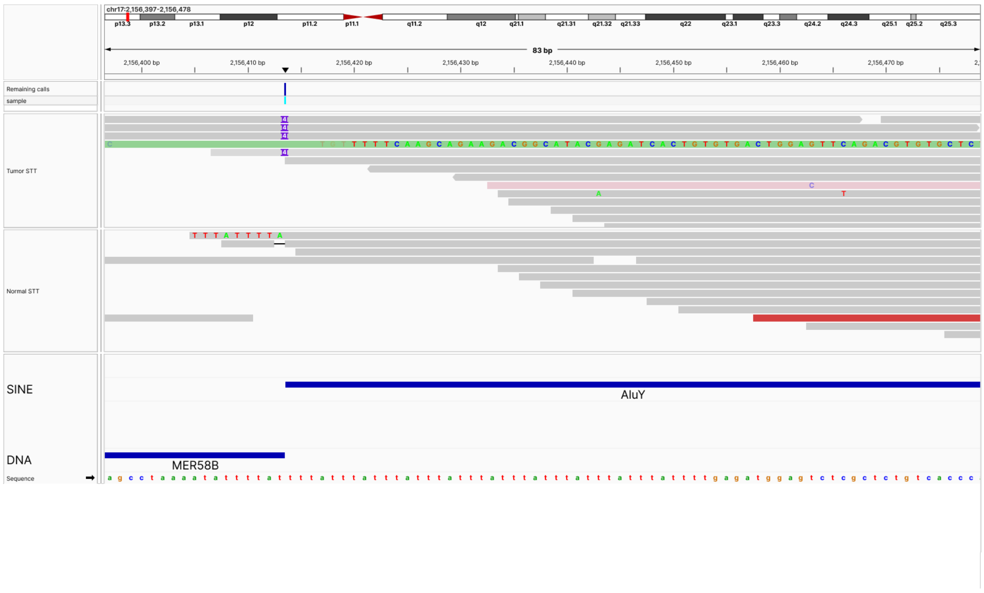


**Fig. S20**: **False positive call (20)**. False positive deletion call predicted by Strelka, likely due to ambiguous read-alignments in this complex region comprising a DNA transposon followed by, followed by a SINE element. This complex repetitive sequence leads to reads mapping with insertions, deletions, and unmapped sequences as soft-clip events. In this case, in the normal sample, there is no evidence of the variant.

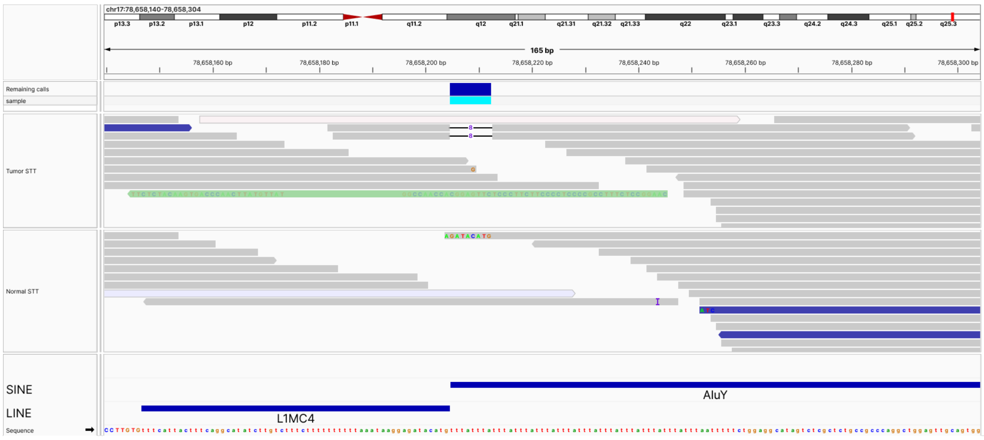


**Fig. S21**: **False positive call (21)**. False positive deletion call predicted by HaplotypeCaller, likely due to ambiguous read-alignments in this complex region comprising LINE and SINE repetitions. This complex repetitive sequence leads to a few reads mapping with deletions and soft-clip events. While the variant caller uses this information to generate a variant call, there is not enough evidence to consider it as a true variation event, instead of a mapping artifact.

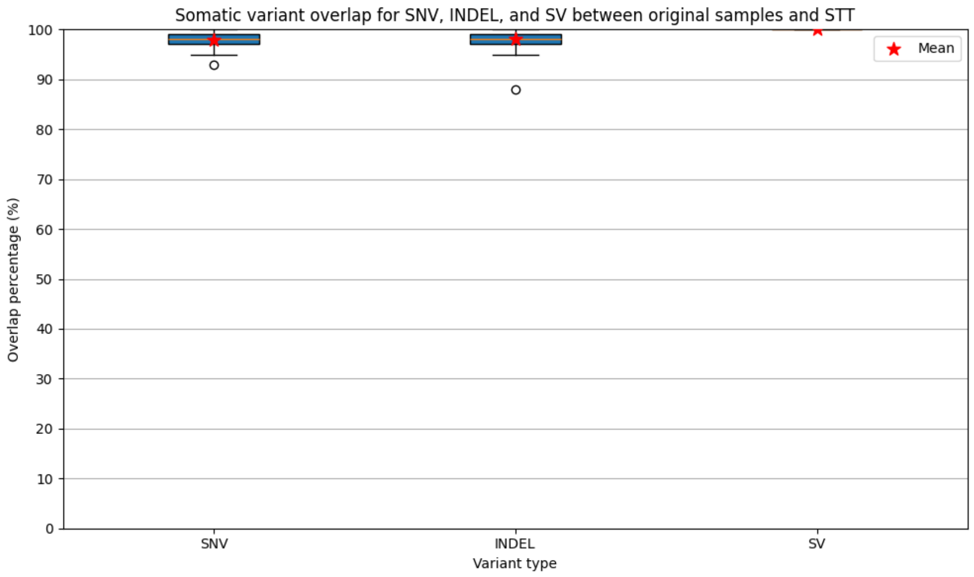


**Fig. S22**: **Somatic variant overlap for SNV, INDEL, and SV between original samples and STT.** Using a previously reported somatic variant identification pipeline (see Methods), from all experimentally validated somatic variants associated with the 47 original tumor samples, their corresponding STTs retained a median of 98±1.4%, 98±2% and 100±4.7% of the original SNVs, indels and SVs, respectively.

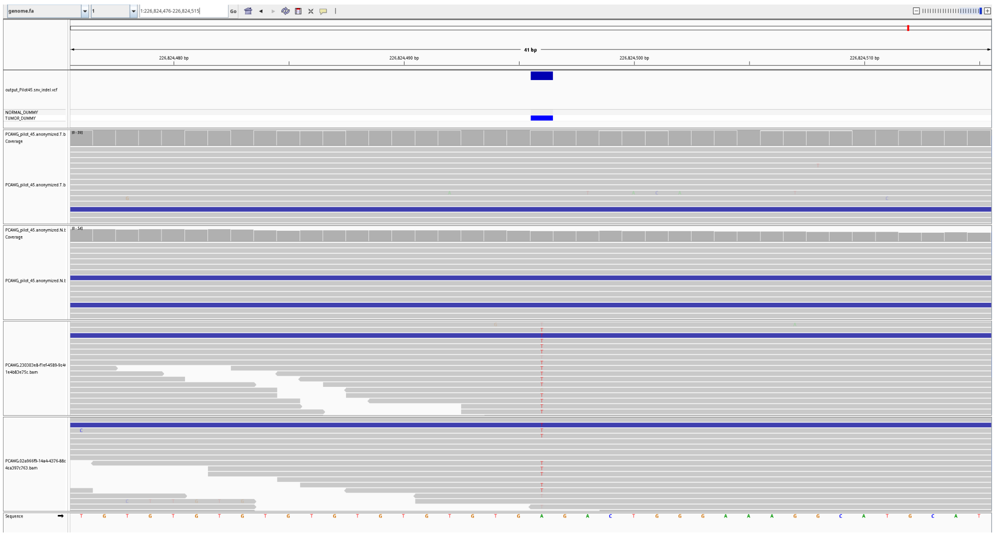

**Fig. S23**: **Correctly removed variant**. This IGV panel showcases the Tumor-Normal STTs and the Tumor-Normal original mappings for a PCAWG-Pilot sample. It can be appreciated how, in the original samples, an alternative T allele is present in multiple reads, both in the tumoral and normal mappings, and was called as somatic. However, this is not the case anymore in the STTs, as it was correctly and completely removed.

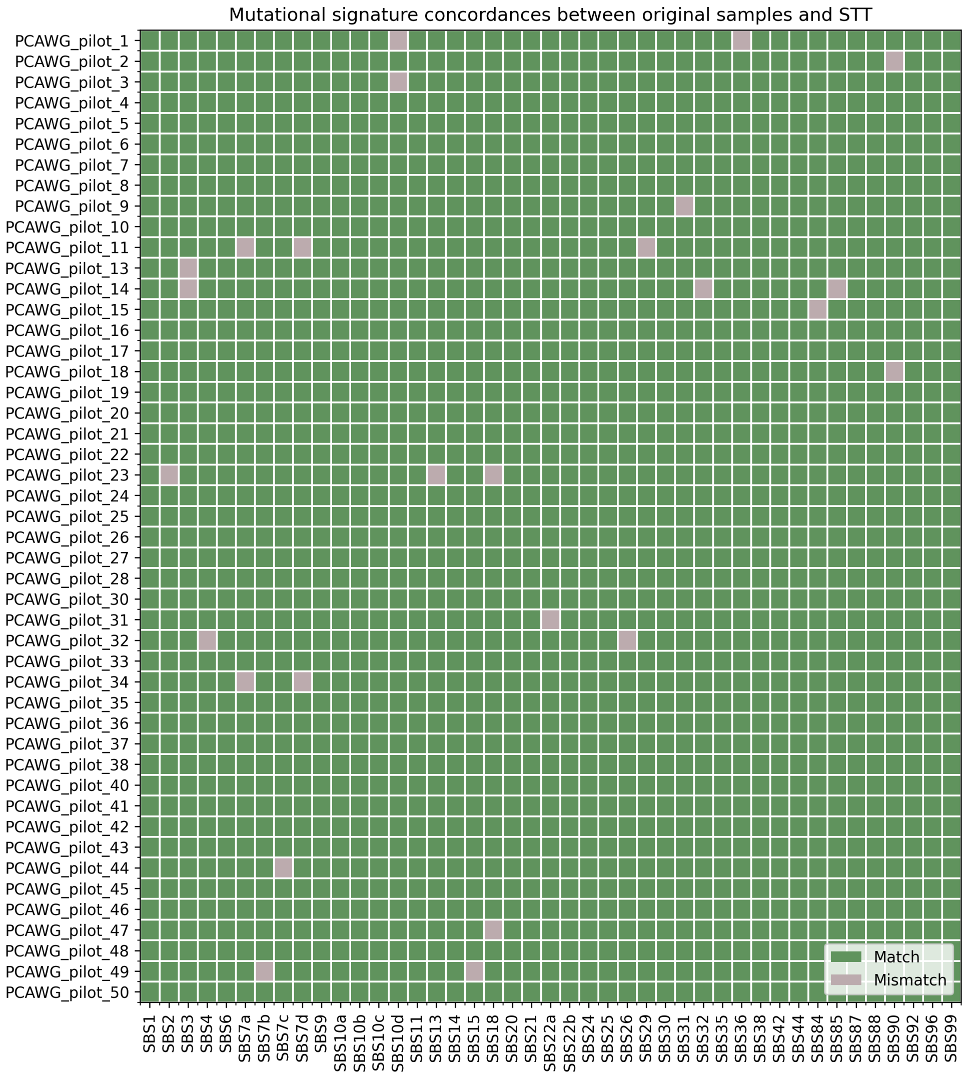


**Fig. S24**: **Concordance of Mutational SBS Signatures in PCAWG-Pilot samples and STTs.** Concordance of 43 known origin mutational signatures between each original PCAWG-Pilot sample and its STT counterpart (see Methods), where matches between them appear in green and mismatches in grey (match if they are present or absent in both samples, and mismatch otherwise). It can be appreciated that there are, approximately, only 1.3% (26) mismatches across all signatures and samples. As detailed in the Methods section, somatic variants were obtained using the curated ONCOLINER discovery pipeline. Then, based on these high-confidence calls, SigProfiler was utilized to determine the presence or absence of each COSMIC SBS signature. Not all unknown origin signatures were included in the analysis.

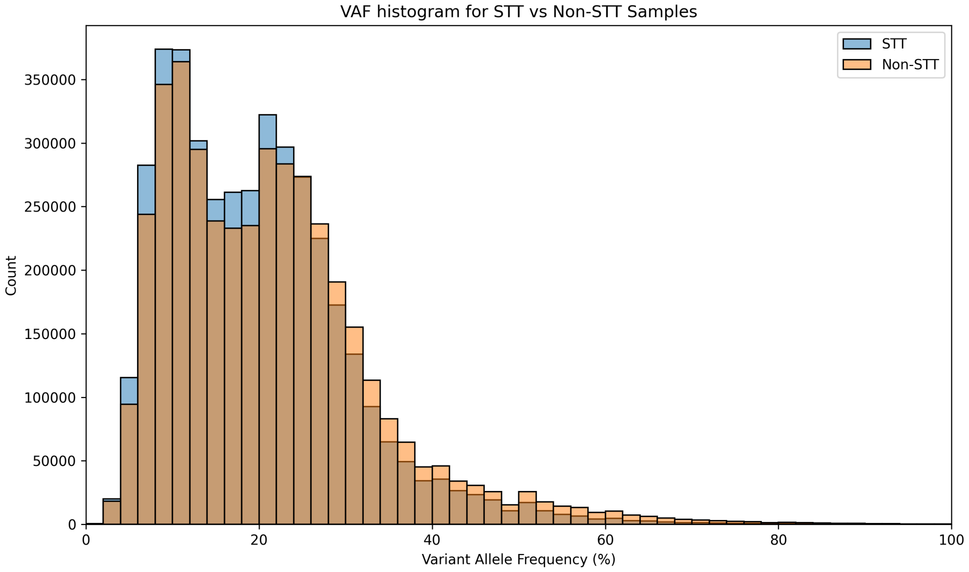


**Fig. S25**: **VAF distributions of somatic variants from PCWAG-Pilot samples and STTs.** Variant Allele Frequency (VAF) distribution from all of the somatic variants called by the ONCOLINER curated pipeline (see Methods), on the original PCAWG-Pilot samples and their STT versions (light brown and blue, respectively). Both VAF distributions are almost identical, with a few differences explained by changes in coverage between the datasets.

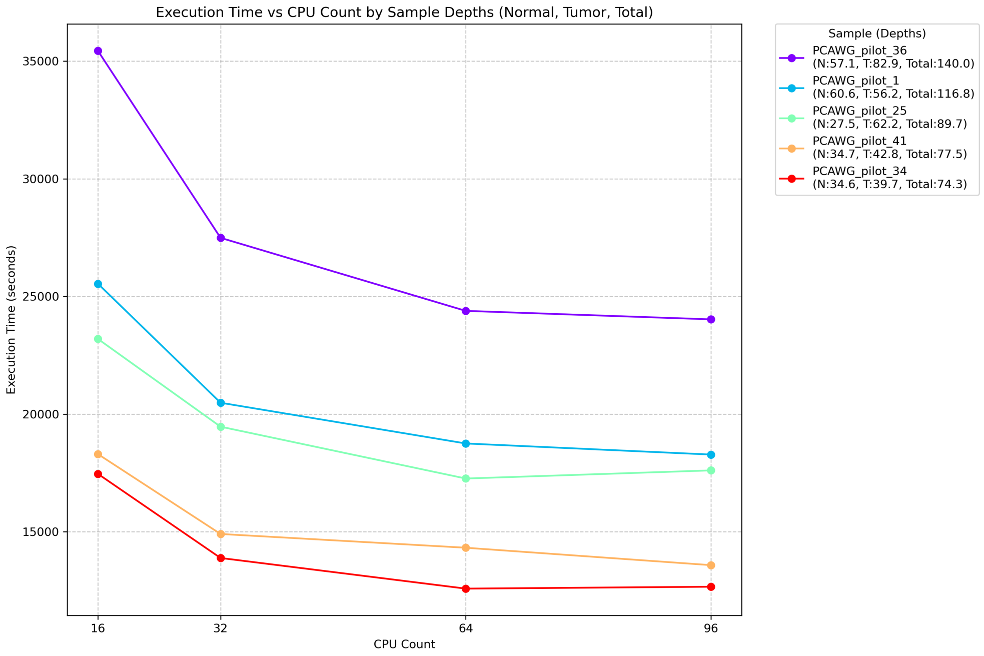


**Fig. S26**: **Performance of the GenomeAnonymizer software under variable CPU counts and datasets with different average depths.** Executing the algorithm in 5 different samples in a single compute node in the MareNostrum 5 HPC (see Methods). The observed trend for each Tumor-Normal Pair shows that the execution time to generate an STT decreases as CPU count increases, with the best performance improvements increasing from 16 to 32 CPUs (1.24 times average speedup). Conversely, the increase from 32 to 64 yielded no significant improvement. The plateau of improvement is because I/O is the main bottleneck. Moreover, the combined depth of each sample pair is the most determining factor for the execution time of the software. Additionally, the required working memory for each experiment was set at around 1.2 times the CPU count (e.g., 22 GB for 16 cores and 36 GB for 32 cores), showing no memory limitation issues.

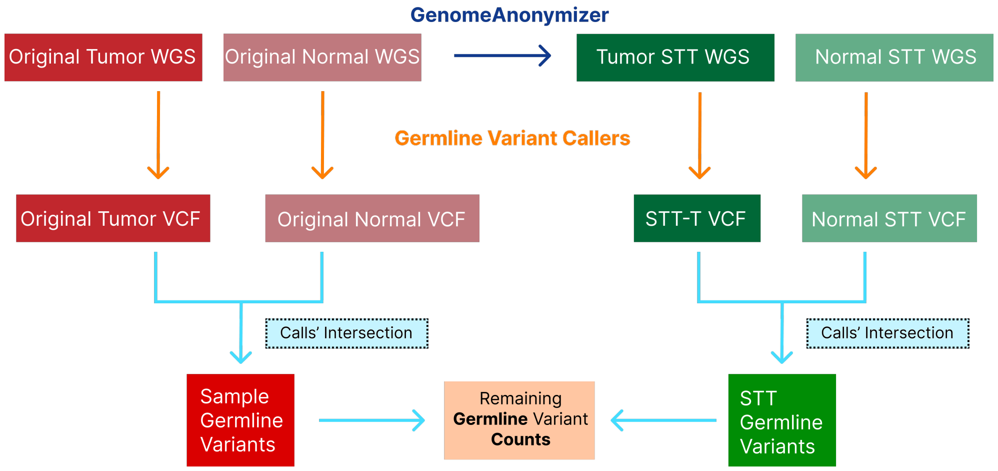


**Fig. S27**: **Strategy to validate the anonymity of the STTs.** To accurately determine if the original germline information remains in the STT, the procedure shown in the figure is executed as follows: 1) The original WGS tumor-normal pairs are anonymized to generate their STT, 2) A selection of 6 state-of-the-art germline variant callers (see Methods) is used to generate variant calls on the tumor and normal mappings, on both the original and STT samples. 3) These calls are intersected between tissues (tumoral and normal) and then between the original and their STT. The final output consists of the intersection of all the variants (if any) called from the original WGS sample and the STT version.

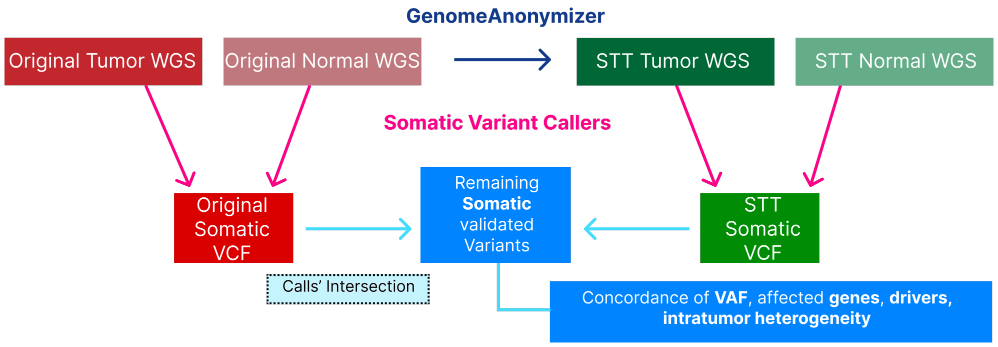


**Fig. S28**: **Strategy to validate conservation of tumor analysis functionality of the STT.** To validate the effective retention of somatic information in the STT of a WGS sample compared to the original sample, the following procedure was devised: 1) An accurate somatic variant discovery pipeline (see Methods) is executed on the original Tumor-Normal paired mappings and the corresponding STT paired mappings. 2) The two resulting VCF variant calls are filtered using the validated variants from the PCAWG-Pilot cohort, leaving only true positive variants. 3) The filtered VCFs from the original sample and the STT are intersected to estimate the fraction of somatic variants that remain in the STT after anonymization. 4) Downstream analyses are performed based on this STT variant call collection, such as assessing the concordance of VAF values between the original and STT samples, retention of functionally relevant variants, mutational signatures, and intratumor heterogeneity.
